# FRET-guided modeling of nucleic acids

**DOI:** 10.1101/2023.08.07.552238

**Authors:** Fabio D. Steffen, Richard A. Cunha, Roland K.O. Sigel, Richard Börner

## Abstract

The functional diversity of RNA is encoded in their innate conformational heterogeneity. The combination of single-molecule spectroscopy and computational modeling offers new, attractive opportunities to map structural transitions within nucleic acid ensembles. Here, we describe a framework to harmonize single-molecule FRET measurements with molecular dynamics simulations and *de novo* structure prediction. Using either all-atom or implicit fluorophore modeling we recreate FRET experiments *in silico*, visualize the underlying structural dynamics and quantify the simulated reaction coordinates. Using multiple accessible-contact volumes (multi-ACV) as a *post-hoc* scoring method for fragment-assembly in Rosetta, we demonstrate that FRET effectively refines *de novo* RNA structure prediction without the need of explicit dye labeling. We benchmark our FRET-assisted modeling approach on double-labeled DNA strands and validate it against an intrinsically dynamic manganese(II)-binding riboswitch. We show that a FRET coordinate describing the assembly of a four-way junction allows our pipeline to recapitulate the global fold of the riboswitch with sub-helical accuracy to the crystal structure. We conclude that computational fluorescence spectroscopy facilitates the interpretability of dynamic structural ensembles and improves the mechanistic understanding of nucleic acid interactions.

**Graphical abstract:** Schematic workflow of integrative FRET modeling using all-atom fluorophores or an accessible-contact volume dye model. All-atom molecular dynamics track the dye coordinate explicitly as part of the simulation while multi-ACV infer mean dye positions *post hoc*.

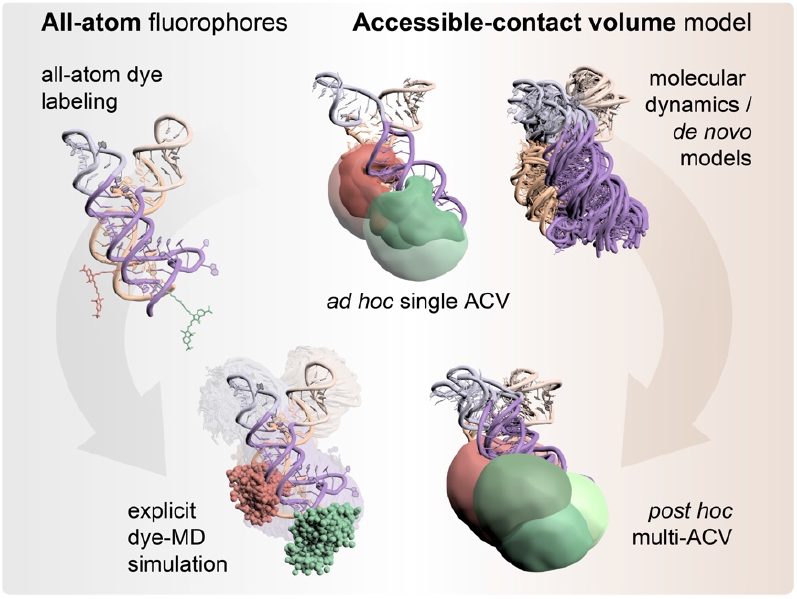

## Introduction

Integrative structural modeling of nucleic acids and proteins has surged in recent years owing to an increasing availability of experimental restraints obtained by various biophysical techniques (1, 2). Such restraints include electron density maps (3, 4), chemical cross-links detected by quantitative mass-spectrometry (5, 6) and distances measured by electron paramagnetic resonance (EPR) with pulsed electron-electron double resonance spectroscopy (PELDOR/DEER) (7, 8) or via Förster resonance energy transfer (FRET) (9–12). In combination with single-molecule detection, FRET lends itself to hybrid modeling as it can resolve short-lived conformational states (13–16) hidden in ensemble-averaged measurements. The high dynamic range of fluorescence spectroscopy therefore ideally complements the spatial resolution of crystallography, nuclear magnetic resonance (NMR) and cryo-electron microscopy (17–20).

Here, we leverage single-molecule FRET for integrated structural modeling of nucleic acids. Previous work in FRET-assisted modeling has focused predominantly on proteins (9–12, 21). Nucleic acids and RNA pose particular challenges with respect to dye labeling (22) and intrinsic domain dynamics (23). With the ribose pucker and six torsion angles in the sugar-phosphate backbone, RNA molecules sample an increased conformational space compared to proteins. Even small RNAs are rarely defined by a single structure but rather exist as dynamic structural ensembles (23, 24). The distribution of these conformers changes in response protein binding (25–27), small-molecule interactions (28, 29) or metal ion coordination (15, 30). Depending on the size and complexity of the RNA, the structures resolved by experimental methods usually represent the most stable conformations. Molecular dynamics simulations address this limitation by computationally sampling trajectories and thus literally fill in the gaps with intermediary states (31).

Furthermore, *de novo* structure prediction has matured to a degree where energetically favorable structures can be computed directly from the nucleic acid sequence, hence bypassing the need for an experimental, structural basis. Most approaches use fragments from libraries of annotated RNA motifs (32, 33) to iteratively assemble larger and more complex architectures. Series of blind prediction challenges on various RNA structures have repeatedly ranked fragment assembly methods amongst the top scoring algorithms for RNA *de novo* modeling (34, 35). These competitions have also highlighted that spatial restraints by chemical footprinting (36) or electron densities (3) help to guide and refine the predicted structures considerably. Incorporating dynamic information, such as FRET-based distance distributions, remains challenging though (10, 11, 17, 37).

To quantitatively connect the readouts of single-molecule FRET measurements to conformational changes, a dye model is required which maps the coordinates of the fluorescence emitters relative to the biomolecule. Here we sketch the blueprints of two forward modeling strategies, both designed to interweave FRET measurements with conformational ensembles. The first approach describes the use of all-atom dyes covalently linked to the biomolecule of interest giving rise to an atomistic view of dye dynamics. The second builds upon our recently introduced implementation of the accessible-contact volume (ACV) in *FRETraj* (38), modeling dye motion implicitly via a geometrical grid search and delineating the spatial boundaries of dye probe (9, 39–41). Either approach aims to localize the donor and acceptor dye to calculate FRET distributions on a structural ensemble *in silico* and subsequently validate the predictions against single-molecule FRET experiments.

We first illustrate concepts, opportunities and pitfalls using a common and well characterized FRET standard, namely a helical DNA fragment labeled at multiple sites with carbocyanine dyes. This molecular-ruler-type benchmark is then complemented by a real-world use case where we model a structurally dynamic riboswitch *de novo*. We demonstrate the feasibility of including experimentally derived FRET constraints as a means to rationally dissect the conformational space, filtering out inconsistent conformers and obtaining a refined ensemble of folded states compatible with FRET experiments.

## Material and Methods

*In silico* pipelines for FRET-assisted structural modeling described herein are logically separated into three steps which involve (1) constructing an DNA/RNA starting structure or *de novo* ensemble, (2) labeling the candidate model with dye probes and (3) predicting FRET and/or dye anisotropy on a series of static structures or a short MD trajectory. Finally, the structural ensemble is validated against single-molecule FRET experiments.

### 1. Nucleic acid structure preparation

DNA double strands were prepared with canonical B-form parameters derived from fiber-diffraction data (42) included in PyMOL 2.3. *De novo* models of the Mn^2+^ riboswitch aptamer were generated using the Fragment Assembly of RNA with Full Atom Refinement (FARFAR2) protocol (32) which is part of the Rosetta modeling suite. The RNA sequence and secondary structure annotation were provided as inputs (Supplementary Methods). The top 500 models (10% of all predicted structures) were ranked by Rosetta energy (33) and used for downstream labeling and FRET predictions.

### 2. *In silico* dye labeling

#### 2.1 All-atom model

All-atom fluorophores were attached *in silico* to C5-amino-modified deoxythymidines on the DNA donor (T19, T23 and T31) and acceptor strand (T31) or to the phosphate at the 5’-ends of a two-stranded Mn^2+^ riboswitch construct using the PyMOL plugin *FRETlabel* (github.com/rna-fretools/fretlabel, Supplementary Fig. 1). The plugin automates the laborious task of manually fusing PDB structures of the dye, linker and nucleic acid of interest. It uses a library of pre-configured dye-linker constructs which reproduce the most common labeling chemistries for nucleic acids (Supplementary Table 1). Force field parameters of the dyes were taken from the AMBERDYES package (43) and expanded to the nucleic acid linkers (Supplementary Methods and Supplementary Fig. 2).

#### 2.2 Accessible-contact volume model

Dye accessible volumes (AV) or accessible-contact volumes (ACV) were computed using a grid search, based on Dijkstra’s algorithm (Supplementary Fig. 3) (39, 44) and implemented by the Python package *FRETraj* (github.com/rna-fretools/fretraj) (38). The AV is the unweighted accessible space that a bioconjugated fluorophore can explore. It is constructed within a rectangular grid spanned around the fluorophore attachment site. A grid node is assigned to the AV given that (i) the node does not clash with the biomolecule surface and (ii) the node is reachable within a distance *L*_linker_ by a walk on the grid starting at the attachment site.

In the ACV model, the AV is additional divided into two separate subvolumes: the contact volume (CV) is defined as a rim around the biomolecule with a diameter *d*_CV_ (set here to match the middle dye radius *R*_2,dye_, Supplementary Table 2) while the free volume (FV) is the complement volume equaling to AV-CV. The CV accounts for the tendency of the dye to interact with the host biomolecule. The fractional occupancy of the contact volume *χ*_CV_ was calibrated by the ratio of residual and fundamental anisotropy

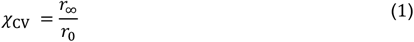

By counting the number of grid nodes in the CV and the FV with weights *ω*_CV_ and *ω*_FV_ respectively, we get

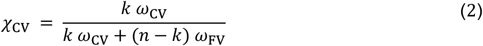

where *n* is the total number of nodes and *k* the number of nodes in the CV. After rearrangement and defining *ω*_FV_ = 1, each node in the CV was assigned a weight

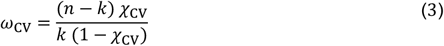

### 3. FRET prediction

Transfer efficiencies between donor and fluorophore were calculated in two ways: (i) with all-atoms dyes or (ii) from a pair of ACVs. In the first case, an inter-dye distance and a *κ*^2^ value were extracted for each frame of an MD trajectory (e.g. every picosecond). The time-dependent *κ*^2^(*t*) was computed from the orientation of the transition dipole vector of the donor and acceptor dye. The instantaneous transfer efficiency *E*(*t*) was then calculated as

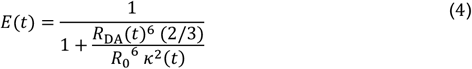

In the second case, an ACV was calculated every 100 ps. The time interval depends on the rotational correlation time and fluorescence lifetime of the dyes. Here, we computed about a dozen ACVs within the fluorescence lifetime (*τ* = 1-1.2 ns) which is sufficient for subsequent sampling of photon events by a Markov process (Supplementary Methods) (45, 46). A mean transfer efficiency Ē_ACV_ was calculated between the donor and acceptor ACV with

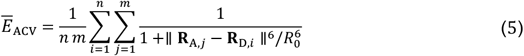

where **R**_D,*i*_ and **R**_A,*j*_ are the coordinate vectors of grid nodes *i* and *j* in the donor and acceptor volume (38, 39, 44).

### 4. Multi-ACV benchmark and riboswitch model selection

The all-atom and multi-ACV based FRET predictions were compared to fully corrected single-molecule FRET measurements. To this end, a thorough photophysical characterization of the donor and acceptor fluorophores was performed by measuring fluorescence lifetime, quantum yields, dynamic anisotropies and burst sizes for the Cy3-Cy5 and Atto550-Atto647N pairs (Supplementary Fig 4). Gamma factors were determined from FRET-stoichiometry (E-S) histograms (47). Computation of multi-ACVs along the MD trajectory reveals the conformational flexibility of the nucleic acid on the timescale of the simulation (1 μs). The predicted energy transfer distribution reflects the distance dynamics without additional broadening by shot noise. To be able to compare the *in silico* FRET histograms against experimental distributions, photon noise was added by simulating fluorescence emission events. Photons were sampled from a Markov chain which defines the time-dependent transfer and fluorescence emission probabilities based on the dye distance *R*_DA_(*t*) calculated between explicit dyes or multi-ACVs. (Fig. 1 and Supplementary Table 3). For all-atom simulations each emitted photon was additionally annotated with a polarization (p = parallel, s = perpendicular) depending on the orientation of the fluorophore dipole.

**Fig. 1.**
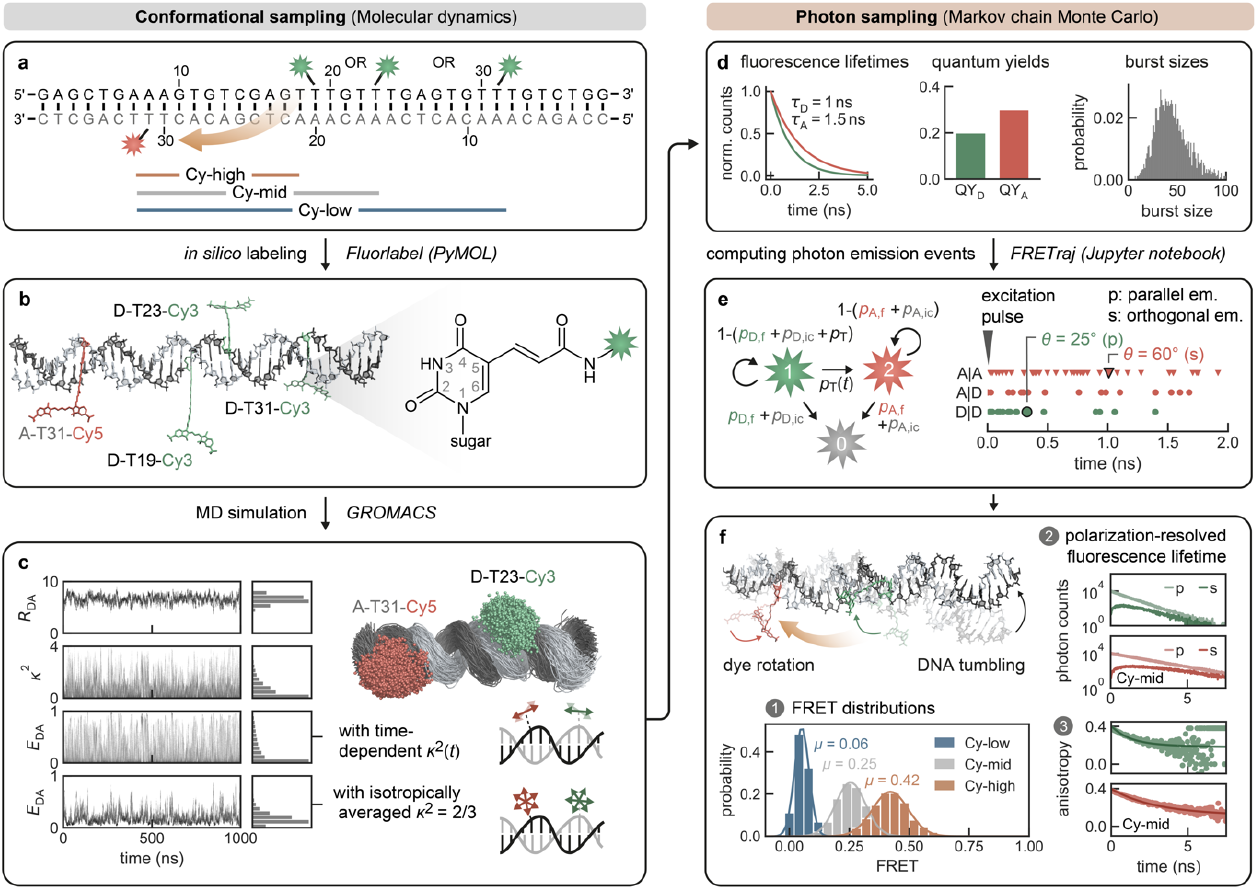
Explicit all-atom fluorescence labeling enable *in silico* FRET prediction on a DNA molecular ruler (49). **a** The double-stranded DNA is labeled at residues T19, T23 or T31 with Cy3 and on the acceptor strand (gray) at T31 with Cy5. The FRET pairs are indicated as Cy-high (12 nt apart), Cy-mid (16 nt apart) and Cy-low (24 nt apart) respectively. **b** *In silico* labeling of the DNA helix via amino-modified deoxythymidines using the PyMOL plugin *FRETlabel*. **c** Structural ensemble from a 1 μs MD simulation showing point clouds of T23-Cy3 and T31-Cy5. Represented are the central carbon atoms of the polymethine chain of Cy3 and Cy5 respectively. The dye distance *R*_DA_, orientation factors *κ*^2^ and instantaneous FRET efficiency *E*_DA_ (with either time-dependent *κ*^2^(*t*) and time-averaged *κ*^2^=2/3) are monitored over the time course of the simulation. **d** Experimentally derived parameters (fluorescence lifetimes, quantum yields and burst sizes) define the relaxation rates in the Markov-chain. **e** Donor and acceptor emission events after donor excitation (D|D and A|D, green and red circles) or by direct excitation of the acceptor (A|A, red triangles) are simulated by Monte-Carlo sampling and are assigned a polarization depending on the angle *θ* between the dye transition dipole at the time points of excitation and emission. **f** Shot-noise limited FRET distribution calculated by averaging over bursts from the 1 μs MD trajectory (all dynamics in the trajectory are fast compared to the burst duration, i.e. there are no slow conformational transitions). The polarization resolved fluorescence lifetime and the derived dynamic anisotropy suggest some interactions of the dyes with the nucleic acid. [double column Fig.]

Riboswitch models were selected by Rosetta energy and filtered by an experimentally informed FRET threshold. The best models were validated against the crystal structure by RMSD and assessed by interaction network fidelity (INF) and deformation index (DI) (48) analogously to quality assessments in RNA puzzles. Additional details are given in the Supplementary Methods.

## Results

### All-atom dye simulations recapitulate single-molecule FRET distributions

Molecular dynamics simulations are a powerful companion approach to structurally interpret single-molecule FRET experiments. To calibrate our *in silico* FRET predictions we first used a well-established FRET standard, a canonical DNA double helix. Specifically, we computationally and experimentally labeled a 38-nt double-stranded DNA at three distinct donor sites and a single acceptor position (Fig. 1a/b). Three separate FRET distributions with the cyanine pair sCy3/sCy5 were measured on a confocal, single-molecule microscope spanning a range between low (Ē_low_ = 0.15), intermediate (Ē_mid_ = 0.39) and high transfer efficiencies (Ē_high_ = 0.58) consistent with a previous multi-laboratory benchmark study (Supplementary Fig. 3 and Supplementary Methods) (49). Next, we simulated 1 µs FRET-MD trajectories for each dye pair to obtain an ensemble of structures and dye positions (Fig. 1c). This point cloud depicts the motion of both the DNA and the fluorophore. From the time-dependent inter-dye distance *R*_DA_(*t*) and orientation factor *κ*^*2*^(*t*) we derived instantaneous transfer efficiencies *E*_DA_(*t*). As this quantity is not accessible experimentally, we recalculated *E*_DA_(*t*) with a time-averaged *κ*^2^=2/3. This reduces the noise in transfer efficiency distribution but does not yet account for dye photophysics. For a more rigorous approximation to the single-molecule experiment, we therefore computed a series of photon bursts incorporating burst sizes, fluorescence lifetimes and dye quantum yields (Fig. 1d/e). The resulting FRET histogram describes a shot noise limited distribution with fast bending dynamics compared to the burst duration (Fig. 1f). The computed anisotropy decays of sCy3 and sCy5 indicate some weak sticking of the fluorophores to the DNA strands consistent with time-correlated single-photon counting (TCSPC) measurements (Fig. 1f and Supplementary Fig. 4) and in accordance with previous results (41, 50–52).

### Accessible-contact volumes are a lightweight model for anisotropic dye distributions

All-atom FRET-MD simulations with fluorophores explicitly included in the force-field are arguably the most precise approach to model dye mobility (Fig. 2a). However, the atomistic precision comes with a few significant downsides: First, explicit dyes and their linkers need to be carefully parameterized for a particular force-field (see Supplementary Methods). Dye packages for the most common force-fields and modeling suites have been published (43, 53, 54) but the list of available fluorophores and adaptors is non-exhaustive, thus often requiring manual modification of PDB structures and topologies. Second, each new dye position typically needs its own simulation. Only if relevant labeling sites were known a priori and independent dyes were not to interact, multiple FRET pairs can be run together. Depending on the size of the biomolecule of interest and the density of labeling sites, unwanted dye interference however often precludes multiplexed simulations.

**Fig. 2.**
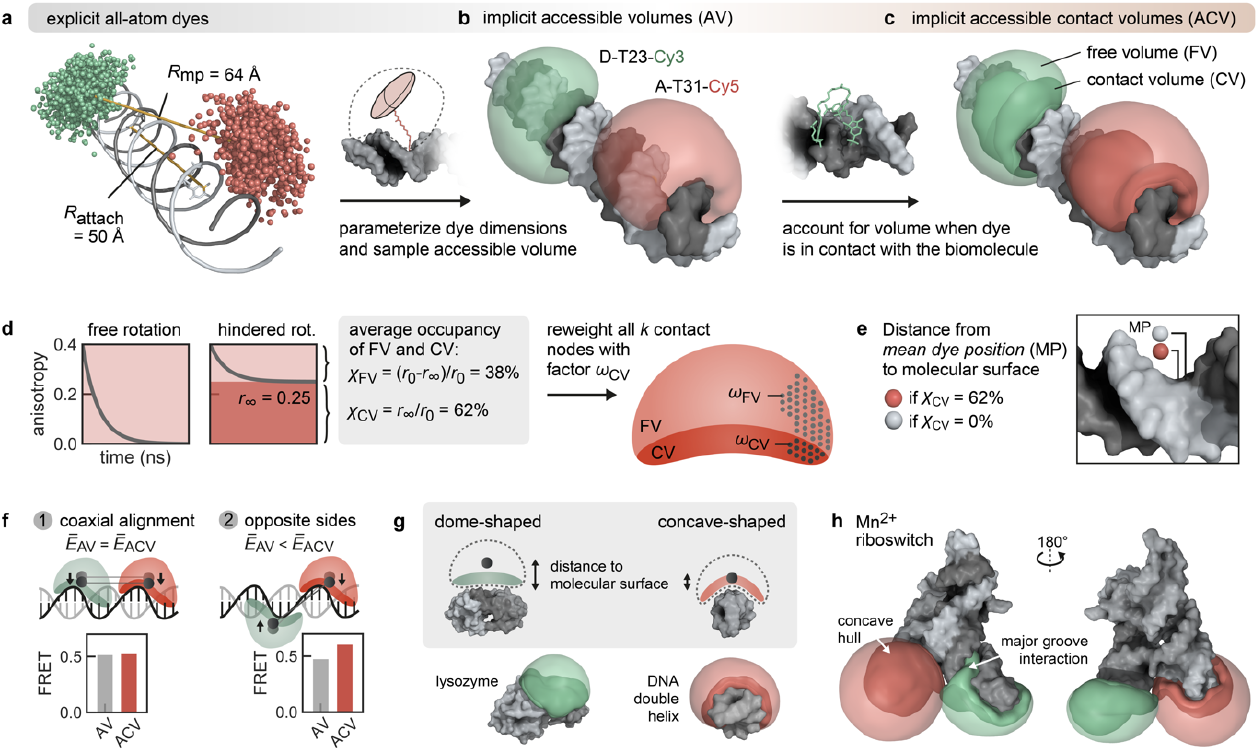
The accessible-contact volume (ACV) represents an anisotropic dye model incorporating free and surface-interacting fluorophores. **a** Comparison of the distance *R*_attach_ and *R*_mp_ from explicit dye simulations on a DNA helix. *R*_attach_ is the distance between the attachment sites (here C5 of dT). *R*_mp_ represents the distance between the center of the dye point cloud sampled in the MD simulation. Because the dyes are coupled to the biomolecule via flexible linkers pointing outwards, it holds that *R*_attach_ < *R*_mp_. **b** The dye point cloud is modeled by accessible volumes (AV) after parameterizing the dye probe as an ellipsoid. (9, 39, 41). **c** The accessible contact volume (ACV) incorporates a rim around the DNA denoted as the contact volume (CV, dark red), which occupied when the dye sticks to the molecular surface. **d** Grid nodes in the CV are reweighted to account for the 62% occupancy of the CV as determined by time-resolved anisotropy (Eq. 1-3). **e** The CV fraction *χ*_CV_ influences the mean dye position (red or white sphere) in the volume. **f** The ACV orientation affects the predicted FRET efficiency Ē_ACV_ only when ACVs are on opposite side of a poly-GC DNA helix (case 2). **g** The biomolecular topology modulates both the ACV shape and the mean position of the dye. The low curvature of globular proteins like lysozyme (PDB: 2lzm) results in a dome-shaped ACV. On the contrary, the ACV wraps around double-stranded DNA with the concave shape shifting the mean position closer to the molecular surface. **h** The Mn^2+^ riboswitch (PDB: 6n2v) features a complex architecture where ACVs in P2 and P4 highlight potential interaction sites for the dyes in the major groove. [double column Fig.]

*Post-hoc* dye models, which can be applied upon an existing MD simulation, are thus an attractive alternative to mitigate the shortcomings of all-atom simulations. Different models have been proposed to address the dye diffusion around an attachment site (9, 39–41, 55). As dyes tend to interact with the host molecule to various degrees, we have previously expanded the widely used accessible volume (AV) model (Fig. 2b) by treating the immediate environment around the biomolecule as a separate phase space with higher weight (Fig. 2c) (38, 41). This reweighting of the accessible volume by the dynamic anisotropy repositions the dye centroid closer to the biomolecule which influences FRET under certain constellations (Fig. 2d/e). In fact, the effect on FRET is expected to be maximal if the ACV clouds are positioned diametrically across the helix and minimal when directly juxtaposed. To quantify the change in FRET by the contact volume, we calculated donor and acceptor ACVs at different sites on a poly-GC DNA strand (Fig. 2f). Inclusion of the contact volume increases the predicted mean FRET value by 0.13 FRET units (Ē_ACV_ = 0.61 vs. Ē_AV_ = 0.48) if the ACVs are on opposite sides of the helical strand but has little to no effect if they are coaxially aligned (Ē_ACV_ = 0.54 vs. Ē_AV_ = 0.52). This is consistent with the outcome of two independent FRET modeling campaigns targeting the proteins atlastin-1 and lysozyme where the ACV model has shown to improve the accuracy of the predicted FRET values compared to the unweighted AV (9, 14).

Not only the relative orientation but also the shape of the dye-accessible volume can affect FRET calculations. The form of the ACV depends on the topology of the biomolecule’s van-der-Waals surface. Globular proteins such as lysozyme, which are devoid of deep cavities, produce ACVs which appear as half domes (Fig. 2g). Helical structures on the other hand lead to concave volumes as the ACV wraps around the spiraling stem. This is exacerbated with folded RNA molecules where intertwined helices expose grooves into which fluorophores may intercalate and get trapped. This is exemplified here by a metal-sensing riboswitch, labeled at the 5’-end and at an internal loop. The ACV is compressed between adjacent stem loops, increasing the fraction of CV nodes within the entire AV and reflecting the increased likelihood for dye-RNA interactions (Fig. 2h). The degree of AV compaction and the relative contribution of the CV was shown to correlate with the extent of fluorescence enhancement (PIFE/NAIFE) (41, 56). Surface interactions of dye and RNA are further enhanced by coordinated metal ions (41) as they bring together distant RNA domains to form tertiary contacts (46, 52, 57) as in the case of the herein studied Mn^2+^ riboswitch. Like with all-atom dye simulations, it is therefore recommended to always visually inspect the resulting ACVs.

### Multi-ACVs predict structural dynamics without explicit all-atom labeling

Having a *post-hoc* dye model at hand that accounts for dye-surface interactions, which are observed to a limited extent even with our FRET standard, we next asked if the ACV model can fully substitute all-atom dyes in an MD simulation. To address this, we calculated multiple ACVs in 100 ps intervals along an MD trajectory of the 38-nt DNA helix (Fig. 3a/b). At each ACV timestep a mean dye distance *R*_DA_ was derived, monitoring DNA bending (58) on a sub-microsecond timescale. The bending angles show a positively skewed normal distribution with a maximum around 13°. Alignment of the DNA backbone produces coalescent volumes (multi-ACVs, Fig. 3c) from which we extracted mean dye positions for each frame (Fig. 3d). The resulting point cloud represents the DNA dynamics broadening the FRET distribution in the single-molecule experiment along with photon noise (Fig. 3e). To incorporate the latter and to correct for quantum yield differences between the dyes, we again simulated photon emission events akin to all-atom dye simulations but here fixing *κ*^2^=2/3 (Fig. 3f). The assumption of isotropic rotational averaging is justified by the observation that mean *κ*^2^ in the explicit dye simulations indeed converges to precisely 0.66 (Supplementary Fig. 5).

**Fig. 3.**
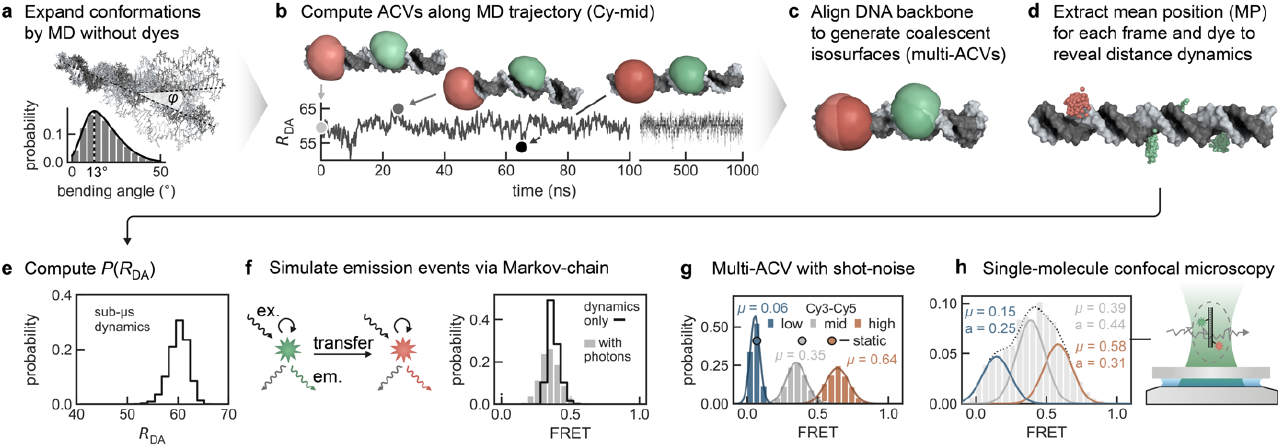
Multi-ACVs simulate FRET-based conformational dynamics without the need of explicit dyes. **a** Bending dynamics of the DNA FRET ruler are sampled by molecular dynamics. **b** Serial ACVs are computed along the trajectory to define an *R*_DA_ time trace with selected snapshots illustrating the bending motions of the DNA. **c** By aligning the DNA, the ACVs coalesce into a merged isosurface (contact volumes were calculated but omitted for clarity in b and c). **d** Dye point clouds representing the mean position of the fluorophore are projected onto the idealized DNA helix. **e** Distribution of *R*_DA_ distances sampled over the converged 1μs MD trajectory. **f** Distances are converted to FRET efficiencies using Eq. 5 (black line). A Markov-chain photon simulation (grey bars) accounts for additional broadening due to shot noise and dye quantum yields. **g/h** Comparison of the predicted FRET histograms from multi-ACV and confocal, single-molecule experiments. Single-ACVs from ideal helices are indicated as spheres. Overall, the mean FRET efficiencies of the three subpopulations in the multiplexed single-molecule experiment are recapitulated well by the simulations. [double column Fig.]

Next, we measured FRET distributions of the three cyanine dye pairs in equal stoichiometries by single-molecule confocal spectroscopy. The measurements were additionally cross-validated by exchanging the cyanine dyes with rhodamine-derived Atto fluorophores, confirming the transfer efficiencies reported in the benchmark study (Supplementary Fig. 4b) (49). A comparison of experiments and simulations shows an overall good agreement of the FRET populations (Fig. 3g/h). Deviations in the relative abundancies are explained by varying dye labeling efficiency and differences in the donor and acceptor noise as a function of FRET. When comparing the position of the FRET peaks we observe that the all-atom dye simulation underestimates the highest FRET pair while the ACV only marginally overestimates the transfer efficiency. This is probably a consequence of a higher degree of dye-stacking interactions in the experiment than suggested by the all-atom MD simulation. The width of the Cy_low_ distribution is broader than predicted by either simulation, potentially due to a higher extent of donor photon noise in the single-molecule experiment.

Altogether, we conclude that the ACV dye model is an appropriate alternative to a full-fledged, all-atom treatment of the fluorophores. ACV calculations provide an intuitive visualization of the dye positions on the biomolecule and help interpret abstract FRET coordinates especially when labeling sites are surface-exposed. Buried attachment sites and nearby cavities pose some more challenges as they contract the ACV, consistent with the concept of nucleic acid induced fluorescence enhancement. We postulate that the implicit ACV model proves particularly useful in the following two scenarios: first, FRET predictions on a single structure can be performed very fast without running an MD simulation first. Second, multi-ACVs can effectively score hundreds of *de novo* modeled candidate structures using experimentally derived FRET restraints.

### FRET selects *de novo* predictions of an intrinsically dynamic riboswitch

Recent RNA puzzle and CASP challenges have demonstrated that current RNA *de novo* prediction algorithms can generate models with remarkable similarity to crystal structures (34, 35, 59). Their scoring metrics still struggle with predicting non-canonical base-pairs and unambiguously ranking top models. Orthogonal assessment methods, particularly measurements at a single-molecule level, may greatly enhance model selection by (in)validating structures with additional constraints. Here, we apply a single FRET coordinate to filter *de novo* modeled riboswitch aptamers consisting of two sets of coaxially stacked stem loops interconnected by a four-way junction (Fig. 4a). Formation of the paperclip-like fold was previously monitored by single-molecule FRET using fluorescent labels at either of the distal helical legs (Fig. 4b) (20). We aimed to reproduce the global H-like architecture of the crystal structure from the nucleotide sequence, its secondary structure annotation and a one-dimensional FRET coordinate (Supplementary Fig. 6a/b).

**Fig. 4.**
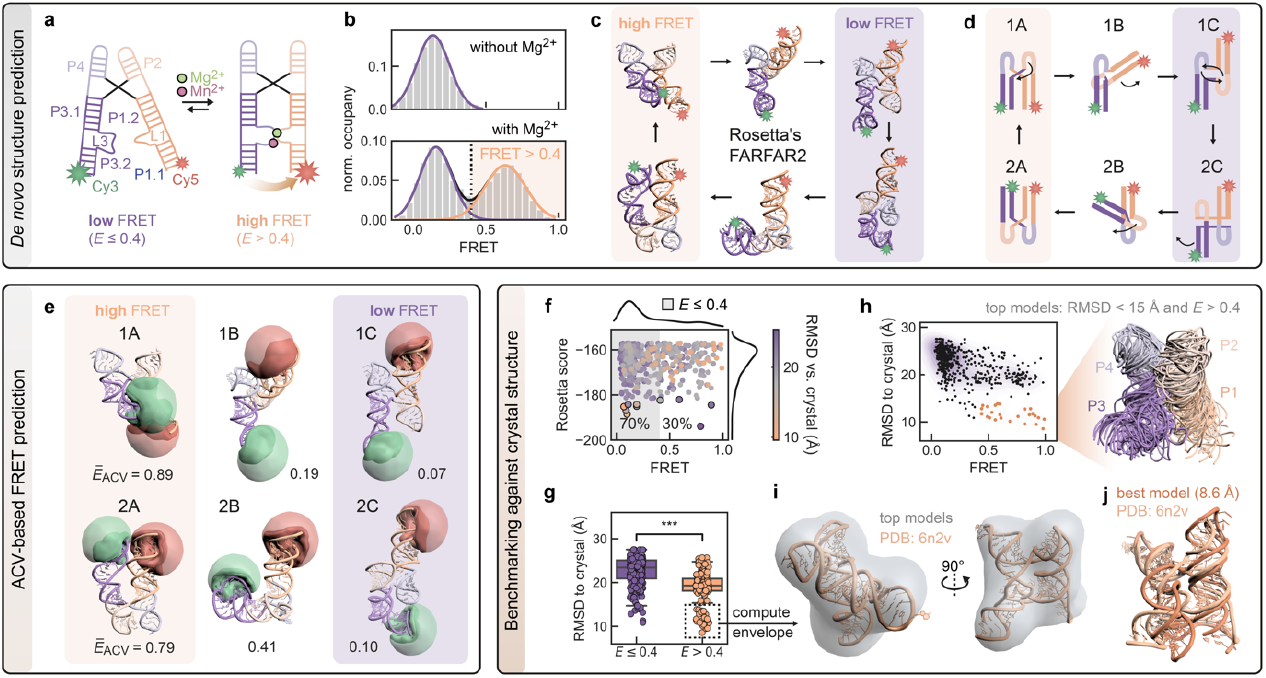
A proof-of-principle FRET-assisted *de novo* modeling pipeline selects candidate structures of a dynamic Mn^2+^ riboswitch compatible with single-molecule FRET experiments. **a** Secondary structure of low and high FRET folding states of the riboswitch mediated by Mg^2+^ and Mn^2+^ binding. **b** FRET histogram illustrating the docking dynamics of the distal helical legs of the riboswitch leading to a high FRET population >0.4. Data from ref. (20). **c** To predict the structure of the folded FRET state *de novo*, 5000 candidate models are generated by Rosetta from the nucleotide sequence (Supplementary Fig. 6). **d** Schematic architectural configurations of the stem-loops P1-P4 categorized by orientation (classes A/B/C) and co-axial stacking (type 1 and type 2, see text). **e** The top 500 models (10% of all models, selected by their Rosetta energy score) encompass candidates of all orientation classes and stacking types. A donor and acceptor ACV is computed for each of these candidates. **f** Next, models were filtered by applying a FRET cutoff at *E* > 0.4 which further narrows down the fraction of candidate structures to 3%. **g** The ensemble of remaining models has a significantly lower mean RMSD to the crystal structure, suggesting that a single FRET coordinate can sort out ill-configured models while retaining native ones. **h** Selection of models with an RMSD < 15 Å to the crystal structure. **i** Overlay of the top models represented as ribbons. **j** Crystal structure aligned into the density map which is computed from the top models. Structural alignment of the best FRET-assisted *de novo* model to the crystal structure (RMSD = 8.6 Å). [double column Fig.]

We first predicted 5000 candidate structures using Rosetta’s fragment assembly of RNA with full atom refinement (FARFAR2) (32). Amongst the top 10% lowest energy models scored by FARFAR2’s energy function (Supplementary Methods), we found a variety of stem loop architectures which were broadly categorizable into three main classes: Class A corresponds to models with slightly twisted but parallel oriented loops, class B represents structural intermediates where one of the helices is rotated by about 90° while class C are fully flipped models with loops pointing in opposite directions. Interestingly, not only the orientation of the loops but also the co-axial stacking of the P1-4 helices can be swapped. This leads to two sets of conformers (type 1 and type 2) each containing the three classes A/B/C and forming a conformational cycle (Fig. 4c/d). It is difficult to assess *per se* which of these models actually do exist on the real folding path towards a native-like structure but the plasticity of four-way junction is still remarkable given that these structures persist after multiple rounds of full-atom refinement.

Next, we predicted FRET efficiencies for each of the 500 top candidates using the ACV dye model. Class A models showed the highest transfer efficiency with values above 0.4, consistent with the fraction of folded riboswitches in the single-molecule histograms, whereas class B and C models are more likely belonging to the ensemble of unfolded or partially folded RNAs (Fig. 4e). Using this FRET threshold, we filter out 70% of the models leaving a subset of structures which on average have a lower RMSD to the crystalized conformer (Fig. 4f/g). It is important to note that a single FRET constraint is not able to pinpoint a single candidate structure that would represent the native fold. However, amongst the models selected by the FRET filter are those candidates with highest similarity to the crystal structure (Fig. 4h). These models recapitulate the global topology of the riboswitch with its co-axially stacked and twisted helices as demonstrated by fitting the crystal structure into the envelope of the aligned structural ensemble (Fig. 4i). The best model approximates the experimental structure with an RMSD of 8.6 Å after all-atom superimposition (Fig. 4j). The metal-binding core with the non-canonical A-minor motif could not be fully reproduced, which is not unexpected given that divalent metal ions mediating core contacts are not considered in the FARFAR2 pipeline (Supplementary Fig. 6c/d). Still, the overall interaction fidelity is reasonably high (INF_all_ = 0.86, INF_WC_ = 0.94, INF_nWC_=0.8), suggesting that most WC and the majority of non-WC contacts which are present in the crystal structure are also formed in the model.

## Discussion

FRET modeling has developed into a versatile tool to resolve short-lived structural ensembles (11, 14, 60–62). Here, we have described two strategies to interface single-molecule FRET measurements with simulations of nucleic acids: the first uses explicit, all-atom fluorophores while the second models dye dynamics implicitly by anisotropy-weighted accessible-contact volumes. Which approach is more appropriate in any given modeling scenario depends on various factors (Table 1).

**Table 1.** Comparison of all-atom fluorophores and the accessible contact volume (ACV) model for FRET predictions.

|  | All-atom fluorophores | Accessible-contact volumes (ACV) |
| --- | --- | --- |
| FRET prediction type | MD simulation | Single PDB structure (crystal or cryoEM structure) or structural ensemble (NMR, MD simulation, *de novo* models) |
| Dye position | Exact position of central dye atom at time $t$ in trajectory | Average position of dye in sterically accessible volume for given structure |
| Computed distance | Exact distance between dye centers at time $t$ in trajectory | Average distance between (multiple) ACVs for given structure or structural ensemble |
| Transition dipole | Yes | No |
| $\kappa^2$ | Time-resolved $\kappa^2(t)$ | Isotropic ($\kappa^2 = 2/3$) |
| Readouts | Instantaneous transfer efficiency $E(t)$ and time-resolved anisotropy $r(t)$ | Mean transfer efficiency $\bar{E}_{ACV}$ |
| Dye labeling during ... | MD setup | Trajectory post-processing |
| Model | Dye / linker force field | Geometrical grid search |
| Implementation | *FRETlabel* | *FRETraj* |
| Photon sampling available (Markov process) | Yes (required) | Yes (optional) |
| Application scenarios | Reproducing single-molecule experiments *in silico* including FRET, fluorescence lifetimes and anisotropy | Finding optimal FRET label positions, filtering *de novo* candidate models and selecting models which are consistent with FRET experiments |
| Computational expenses <sup>1</sup> | Fluorophore sampling (100-200 ns trajectory): hours - days depending on infrastructure | (Multi)-ACV calculation: seconds - minutes on personal computer |

<sup>1</sup> Estimated computational time for sampling the accessible space of the dye around the biomolecule relevant to the choice of the dye model. Time requirement for extensive conformational sampling of biomolecule is system-dependent (size of biomolecule and the water box) and thus not specified here.

All-atom MD are considered the de facto gold-standard for simulation of FRET experiments. Tracking the precise location of the fluorophore makes both translational and rotational dye components available to compute experimental properties including fluorescence anisotropy. Knowledge about the inhomogeneity of a dye’s orientational distribution was factored into the design of the ACV model. Through a linear combination of reweighted subvolumes delineating the accessible space of freely rotating and hindered dyes, we achieve more accurate FRET predictions than with the traditional AV model. Multi-ACVs now open up new avenues to high-throughput *in silico* screening with a plethora of applications for *de novo* modeling, labeling site selection or post-processing of MD trajectories.

Compared to all-atom simulations, ACV screens are straightforward to setup, fast to run and easy to visualize. Being equally applicable to both single structures and ensembles, amenable to proteins and nucleic acids alike, the potential of multi-ACVs lies in the ability to make FRET coordinates and trajectories structurally explainable. For instance, ACV-based FRET predictions can rapidly inform about sensitive donor-acceptor sites to be followed-up experimentally. Another key advantage of *post-hoc* dye models is their applicability to existing or published datasets such as micro-millisecond long MD trajectories without predefined labeling sites. Here, we have carved out important factors that influence ACV-based FRET predictions. Most notably, the shape and orientation of the ACV are modulated by the global and local topology of the biomolecule including the presence of grooves or exposed nucleotides stretches, conditionally affecting FRET.

Finally, we have highlighted present and future challenges in FRET-assisted modeling. The *de novo* prediction of a metal-binding riboswitch nicely illustrates the quest for the proverbial hairpin in the unstructured haystack. Current state-of-the-art structure prediction workflows are able to sample native-like folds amongst many suboptimal conformers. Validating true hits with high confidence is still a major difficulty though and prompts inclusion of complementary biophysical methods. Using FRET as a reaction coordinate, we here dissected the conformational ensemble of the riboswitch and were able to effectively narrow down the list of structural candidates. To increase the specificity of FRET in pinpointing the most native-like models additional, orthogonal distances would need to be sampled experimentally. In the process of building such minimally redundant networks of donor-acceptor pairs (9, 12, 14) ACV screens are instrumental in figuring out which labeling sites to prioritize. Future studies will address how to include the most informative FRET coordinates as active restraints into integrative FRET modeling pipelines of nucleic acids to not only select but also direct conformers along their folding route.

## Acknowledgements

The authors thank Felix Erichson and Besim Fazliji for software testing and feedback. Financial support from the Swiss National Science Foundation (200020_165868 and 200020_192153 to R.K.O.S), the University of Zurich (to R.K.O.S.) and the UZH Forschungskredit (FK-17-098 to F.D.S., FK-73521-10-01 to R.C. and FK-14-096 / 15-095 to R.B.) is kindly acknowledged.

## Contributions

F.D.S. and R.B. designed the research. F.D.S. performed and analyzed single-molecule measurements and molecular dynamics simulations. R.A.C. ran MD simulations. R.B and R.K.O.S supervised the project. F.D.S and R.B wrote the manuscript with contributions from all authors.

## Data availability

The source code of *FRETraj* (prediction of ACVs), *FRETlabel* (labeling of nucleic acids with all-atom fluorophores) and *LifeFit* (reconvolution-based fitting of eTCSPC data) is available at https://github.com/RNA-FRETools/ and is documented at https://rna-fretools.github.io/. The PyMOL plugins are available as Docker containers with bundled open-source PyMOL. The API of *FRETraj* is documented in Jupyter notebooks with step-wise tutorials and examples of RNA structures presented herein. Protocols for dye-linker parameterization with *FRETlabel* are available at https://github.com/RNA-FRETools/fretlabel. The source code for modeling the manganese(II) riboswitch with Rosetta and FARFAR2 is hosted at https://github.com/RNA-FRETools/rosettascripts/. De novo models from the integrative FRET-guided structure prediction trial of the manganese(II) riboswitch can be found at https://github.com/RNA-FRETools/Steffen_FRETguided_modeling_2023.git. Additional source material is available from the corresponding author upon request.

