## Supporting Information for "FRET-guided modeling of nucleic acids"

### Supplementary Methods

#### *In vitro* nucleic acid preparation and dye labeling

DNA oligonucleotides were purchased with C5-amino-modified deoxythymidines at three donor (T19, T23, T31) and one acceptor (T31) label site. The dyes were coupled in-house to the DNA strand using N-hydroxysuccinimide chemistry. In brief, 5 µL of concentrated DNA (1 mM) were mixed with 3 µL of dye (few crystals of NHS-dye ester dissolved in 12 µL DMSO, >10x excess of dye) in 2 µL of potassium phosphate buffer (500mM, pH 8.7) and stirred at 37 °C and 400 rpm for 3 h (1). The DNA was precipitated overnight at -20°C in 50 µL (2.5 eq) of ice-cold EtOH supplemented with 10 µL KCl (1M). The pellet was washed 3x with 70% EtOH, resuspended in water. Excess salts and phosphate buffer were removed by filtering over a VivaSpin (5000 MWCO) and purity was checked by analytical HPLC.

#### Ensemble time-correlated single photon counting (eTCSPC)

Fluorescence lifetime and anisotropy decays were recorded on a Fluorolog FL3-222 (Horiba) spectrophotometer equipped with a double-grating emission monochromator and a PPD-900 photon counting detector. Samples were excited at 504 nm (DD-510L, Horiba) and 633 nm (DD-635L, Horiba) with a repetition rate of 10 MHz. Decays were fitted to a series of exponentials by iterative reconvolution with the instrument response function (IRF) using a custom written Python package named *LifeFit* (https://github.com/RNA-FRETools/Lifefit). The dynamic anisotropy was calculated from the polarization resolved decays

$$r(t) = \frac{I_{\parallel}(t) - G\,I_{\perp}(t)}{I_{\parallel}(t) + 2G\,I_{\perp}(t)} \tag{1}$$

and analyzed with a local-global rotation model (2, 3)

$$r(t) = \left[(r_0 - r_\infty)\,e^{-t/\tau_{r,loc}} + r_\infty\right] e^{-t/\tau_{r,glob}}\,. \tag{2}$$

#### Single-molecule confocal experiments

DNA double strands were imaged free in solution in the presence of 100 mM KCl and 20 mM $MgCl_2$, buffered in 50 mM Tris-HCl at pH 7.5. Single-molecule measurements were performed on an upright MicroTime 100 confocal microscope (Picoquant) with pulsed interleaved excitation (PIE) (4) at 518 nm and 640 nm (LDH-D-C-640 and LDH-D-C-520, Picoquant) and a repetition rate of 40 MHz. The instrument was equipped with a UPlanSApo, 60x, NA=1.20 water-immersion objective (Olympus) and a TimeHarp 260 TCSPC card (Picoquant) (5). Photons were spectrally separated (Semrock FF01-582/64 and Chroma H690/70m filters), directed through fiber optics (pinhole diameter: 50 µm) onto two SPCM-AQRH avalanche photodiodes (Excelitas). The data was analyzed with the Matlab software package *PAM* (6). A sliding time-window with two-color, all photon burst search (APBS) identified bursts with a total count of at least 100 photons ($N_D$ + $N_A$) (7). Double labeled molecules were selected by applying an intensity threshold of red photons after acceptor excitation ($F_{A|A}$ > 50) to remove the donor-only population and a stoichiometry limit $S$ > 0.2 to eliminate any acceptor-only species (8). FRET histograms were corrected for donor leakage into the acceptor channel ($\alpha$ = 0.1 of donor signal), direct excitation of the acceptor ($\delta$ = 0.02 of acceptor signal) as well as differences in detection and quantum efficiency of the two dyes ($\gamma$ = 0.7) by non-linear regression in the 2D FRET-stoichiometry (*E*-*S*) histogram (9, 10).

#### Derivation of fluorophore and linker parameters for *in silico* labeling with *FRETlabel*

FRET labels were composed of three parts: (i) a fluorescent moiety with a delocalized π-electron system, (ii) a flexible linker and (iii) a functional group for conjugation with the biomolecule of interest. We constructed and parameterized suitable adapters for the most common nucleic acid labeling strategies (11). Bonded and non-bonded parameters for the fluorophores were taken from the AMBER-DYES package (12, 13). Parameters for the linker atoms were derived by analogy to similar terms in the AMBER force field using the Antechamber Python Parser interface (Acpype) (14). Linker geometries were optimized with GAUSSIAN 09 (15) at the B3LYP/6-31G* level of theory and followed by electrostatic potential (ESP) calculations with Hartree-Fock. Partial charges were calculated by two-stage restrained electrostatic potential (RESP) (16) fitting, as implemented in Antechamber (17). For this purpose, appropriate capping groups and group constraints were introduced, mimicking the partial charge distribution in the linker when ligated to the dyes and nucleotides (Supplementary Fig. 2). Step-by-step protocols for linker building, RESP fitting and force field parameterization can be found at github.com/RNA-FRETools/fretlabel. Dyes and linkers were fused to the RNA at the desired position using the PyMOL plugin named *FRETlabel*. Through a graphical user interface (GUI) *FRETlabel* exposes ready-to-use dye-linker adaptors for 3'/5'-end labeling as well as internal labeling at deoxythymidines or etheno-bridged adenosines and cytidines (Supplementary Fig. 1).

#### Molecular dynamics simulations

Molecular dynamics simulations were run on Piz Daint (Swiss National Supercomputing center CSCS, Lugano) with Gromacs 2019.4 and 2020.3 (18) using the AMBER force field (19) with parmbsc0 (20) and χOL3 (21, 22) corrections for RNA and ε/ζOL1 (23), χOL4 (24) and βOL1 (25) corrections (together named OL15) for DNA. Ion parameters for $K^+$ and $Cl^-$ are taken from Joung and Cheatam (26). The fluorophore labeled RNA was solvated in a dodecahedral water box with TIP4P-ew (27), charge neutralized with KCl and equilibrated at 298 K and 1 bar with the velocity rescale thermostat and the Berendsen barostat for 10 ns. Bonded interactions were constrained by the LINCS algorithm (28) using an integration time step of 2 fs while non-bonded interactions were treated with the Verlet scheme and a cut-off of 1.4 nm. Long-range electrostatics were computed with the particle mesh Ewald method (29). Perdiodic boundary correction (PBC) was performed with Plumed 2.6.0 (30) reconstituting and translating molecules to align the center of mass. Only a translational (`TYPE=SIMPLE`) but no roto-translational fit (`TYPE=OPTIMAL`) was performed to preserve the global rotational component of the biomolecule for subsequent anisotropy calculations.

```
plumed driver --plumed plumed_pbc.dat --mf_xtc traj.xtc
```

where the configuration file `plumed_pbc.dat` contained

```
nucleic_acid: GROUP NDX_FILE=nucleic_acid.ndx NDX_GROUP=nucleic_acid
WHOLEMOLECULES ENTITY0=dna
FIT_TO_TEMPLATE STRIDE=1 REFERENCE=reference.pdb TYPE=SIMPLE
DUMPATOMS FILE=traj_pbc_translation.xtc ATOMS=nucleic_acid STRIDE=1
```

Distances between the central carbon atoms in the polymethine chain of the donor and acceptor fluorophore were calculated every picosecond. Likewise, the time-dependent orientation factor $\kappa^2(t)$ was computed for each snapshot using `gmx dyecoupl` (31)

$$\kappa^2(t) = (\cos\theta_\mathrm{T} - 3\cos\theta_\mathrm{D}\cos\theta_\mathrm{A})^2 \tag{3}$$

where $\theta_\mathrm{D}$ and $\theta_\mathrm{A}$ are angles between the vector connecting the central carbons and the transition dipole of donor or acceptor respectively, and $\theta_\mathrm{T}$ is the angle between the two transition dipoles (2).

The DNA bending angle $\varphi_\mathrm{bend}$ was calculated using Plumed 2.6.0 as the deviation from the helical axis defined by atoms DC1(N4) – DA19(N6) – DC38(N4)

$$\varphi_\mathrm{bend} = 180 - \angle_{\mathrm{DC1(N4)-DA19(N6)-DC38(N4)}} \tag{4}$$

**Simulation of photon emission events**

Photons were sampled from a Markov chain illustrated in Fig. 1e and characterized by the photon emission rates $k_\mathrm{D,f}$ and $k_\mathrm{A,f}$, the internal conversion rates $k_\mathrm{D,i}$ and $k_\mathrm{A,i}$ and the time-dependent transfer rate $k_\mathrm{T}(t)$. The transition probabilities between the ground state, the donor-excited state and the acceptor-excited state are given as

$$P = \begin{pmatrix} 0 & 1 & 0 \\ p_{D,f} + p_{D,ic} & 1-(p_{D,f}+p_{D,ic}+p_T) & p_T \\ p_{A,f}+p_{A,ic} & 0 & 1-(p_{A,f}+p_{A,ic}) \end{pmatrix} \tag{5}$$

The transfer rate was updated at each time-step of the Monte-Carlo simulation by the inter-dye distance $R_\mathrm{DA}(t)$ and the orientation factor $\kappa^2(t)$ of the underlying MD trajectory

$$k_\mathrm{T}(t) = (k_\mathrm{D,f} + k_\mathrm{D,ic})\left(\frac{R_0'(t)}{R_\mathrm{DA}(t)}\right)^6 \tag{6}$$

with
$$k_\mathrm{D,f} = \frac{Q_\mathrm{D}}{\tau_\mathrm{D}}, \quad k_\mathrm{D,ic} = \frac{1-Q_\mathrm{D}}{\tau_\mathrm{D}} \tag{7}$$

and
$$R_0'(t)^6 = \left(\frac{R_0{}^6}{2/3}\right)\kappa^2(t) \tag{8}$$

where $Q_\mathrm{D}$ and $\tau_\mathrm{D}$ are the fluorescence quantum yield and lifetime of the donor dye (31, 32). $R_0$ is the time-independent Förster radius (54 Å, for Cy3-Cy5 assuming isotropic rotation) while $R_0'(t)$ is time-dependent and is a function of the instantaneous $\kappa^2(t)$. For ACV simulations isotropic rotation of the dye was assumed ($\kappa^2$=2/3).

Burst sizes were drawn from an experimental, single-molecule distribution $P(N_\mathrm{D} + N_\mathrm{A})$ or an analytical power law distribution

$$P(x) = x^\lambda \quad \text{with } x \in \mathbb{N} \text{ and } N_\mathrm{T,min} < x < N_\mathrm{T,max} \tag{9}$$

where $x$ is the burst size, $\lambda$ an empirical parameter to approximate the experimental distribution and $N_{T,min}$ and $N_{T,max}$ lower and upper size cutoffs (31).
For each burst, photons were accumulated until the total photon count $N_T = N_D + N_A$ had reached the current burst size.

Each burst can be computed from a Markov process involving (i) only one MD trajectory or (ii) all trajectories as described previously (31, 32). The first case reflects a situation where interconversion between conformation is slow compared to the duration of the burst (i.e. each photon burst averages only over fast dye motions within one trajectory). In the second case, the entire trajectory ensemble is represented in one burst (i.e. all conformational dynamics of dyes and biomolecule are averaged in one burst). In either way, a transfer efficiency $E$ is computed for each burst

$$E = \frac{N_{A,\mathrm{tot}}}{N_{A,\mathrm{tot}} + N_{D,\mathrm{tot}}} \quad (10)$$

where $N_{A,tot} = N_A/Q_A$ and $N_{D,tot} = N_D/Q_D$. The resulting histograms can be compared to a fully $\gamma$-corrected FRET experiment. Here we calculated FRET distributions for each donor-acceptor DNA strand pair separately to get their respective mean FRET efficiency. Additionally, we reproduced the multiplexed FRET experiment by combining all three 1 ns simulations (Cy-low, Cy-mid and Cy-high) using the “one-trajectory” averaging regime described above.

By calculating photon emission events, we account for shot noise broadening observed in experimental FRET histograms. The variance of the resulting distribution is therefore the sum of the shot-noise contribution and the variance induced by structural dynamics ($\sigma_{dyn}$) (33)

$$\sigma^2_{\mathrm{sn+dyn}} = \frac{\langle E\rangle_{\mathrm{exp}}(1-\langle E\rangle_{\mathrm{exp}})}{N_\mathrm{A} + N_\mathrm{D}} + \sigma^2_{\mathrm{dyn}}\,. \quad (11)$$

There are two notable limitations to the burst generation method outline herein. Firstly, donor and acceptor brightness are assumed to be constant and independent of the instantaneous transfer efficiency. Secondly, the photon simulation does not include background noise and therefore should be compared to background corrected FRET experiments.

For dynamic anisotropy decays, the photon polarization probability was calculated as

$$P(\mathrm{p}) = \frac{2\cos^2(\theta)}{\cos^2(\theta)+1} \quad \text{and} \quad P(\mathrm{s}) = 1 - P(\mathrm{p}) \quad (12)$$

where $\theta$ is the angle between the transition dipole at the time points of excitation and emission respectively. Apart from depolarization due to rotational diffusion, the angular displacement $\beta$ between the absorption and emission dipole as well as photoselection (prefactor: 2/5) also decrease the anisotropy (32). The time-resolved anisotropy $r(t)$ is thus expressed as the product of three terms

$$r(t) = \frac{2}{5}\left(\frac{3\cos^2(\beta)-1}{2}\right)\left(\frac{I_p(t)-I_s(t)}{I_p(t)+2I_s(t)}\right) \quad (13)$$

where $I_p$ and $I_s$ are the intensities of the polarization resolved lifetime decays.

**Riboswitch structure prediction with Rosetta**

For Rosetta modeling of the $Mn^{2+}$ sensing *yybP-ykoY* riboswitch aptamer from *Xanthomonas oryzae* (34) we used the FARFAR2 protocol (35) which is part of Rosetta 3.12 (https://www.rosettacommons.org/) and includes improved base-pair sampling as well as an updated fragment library (36) and scoring function ("res4") (37). Models were calculated with `rna_denovo`, executed on multiple cores in parallel:

```
rna_denovo.linuxgccrelease -nstruct 1000 -fasta mn_riboswitch.fasta
-secstruct_file mn_riboswitch_secstruct.txt -silent mn_riboswitch.out
-minimize_rna true -cycles 20000
```

As inputs only the FASTA sequence of the FRET construct (34) and its secondary structure annotation were provided:

```
>mn_riboswitch A:1-57 B:1-48
auccuuggggaguagccugcuuucuucggaaagcgccuguaucaacauacucggcua,uagccguggu
gcaggcaacggcgaaagccgucuggcgagaccagggau

((((((((......(((.((((((....))))))((((((((((.......((((((,))))))))))
)))))).(((((....)))))..)))....))))))))
```

Supplementary Fig. 6a shows an alignment of the FRET construct and the crystal structure (PDB entry 6n2v).
The top-500 lowest energy models out of a total of 5000 structures calculated by FARFAR2 were extracted with Rosetta's `extract_pdbs`

```
grep "^SCORE:" mn_riboswitch.out | grep -v description | sort -nk2 |
head -n 500 | awk '{print $NF ": " $2}' | tee top_models.txt

extract_pdbs.linuxgccrelease -in:file:silent mn_riboswitch.out -tags
`cat top_models.txt | cut -f1 -d':'`
```

The extracted PDBs were merged into a single file `mn_riboswitch.pdb`

```
for pdb in `cat top_models.txt | cut -f1 -d':'`; do
  echo MODEL $i >> mn_riboswitch.pdb
  echo TITLE "$pdb" >> mn_riboswitch.pdb
  cat "$pdb".pdb >> mn_riboswitch.pdb
  echo -e "ENDMDL\n\n" >> mn_riboswitch.pdb
  i=$(($i+1))
done
```

Sequence agnostic RMSD was calculated for all top-500 RNA models after superimposition to the crystal structure using PyMOL's `align` function

```
rmsd = []
for in in range(1,501):
  rmsd[i]=cmd.align(f"mn_riboswitch and state {i}", "6n2v_conformer",
  mobile_state=i, cycles=0, transform=0)[0]
```

To calculate metrics relying on exact sequence identity, the sequence of the FRET model was threaded onto the crystal structure (i.e. mutating residues of the crystal structure to match the nucleotides of the FRET construct) using Rosetta's `rna_thread`

```
rna_thread -in:file:fasta alignment.fasta -s 6n2v_conformer.pdb -o
6n2v_with_FRETsequence.pdb
```

where `alignment.fasta` contained the following sequence alignment:

```
>mn_riboswitch A:2-58 B:1-15 B:18-19 B:22-27 B:30-47
uccuuggggaguagccugcuuuc-uuc-ggaaagcgccuguaucaacauacucggcua,
uagccguggugcaggaccgaaaggucuggcgagaccaggga
```

```
>6n2v_conformer1.pdb  B:1-99
ggcuuggggaguagccugcuuucggaaacgaaagcgccuguaucaacauacucggcgaaagccguggu
gcaggaccgaaaggucuggcgagaccaggcc
```

The threaded target structure was renumbered with `pdb_resi_renumber` from *rosettascripts* (https://github.com/RNA-FRETools/rosettascripts) to conform to the FRET construct sequence in Supplementary Fig. 6a/b.

```
pdb_resi_renumber -pdb 6n2v_with_FRETsequence.pdb -e 'A:1-97>A:2-
57,B:1-15,B:18-19,B:22-27,B:30-47'
```

RMSD and Interaction Network Fidelity (INF) metrics were calculated over residues A:2-24, A:28-57, B:1-15 and B:30-47 using `rna_tools` (38) akin to quality assessment in RNA puzzles

```
rna_calc_rmsd.py -t 6n2v_with_FRETsequence_renum.pdb --target-selection
A:2-24+A:28-57+B:1-15+B:30-47 --model-selection A:2-24+A:28-57+B:1-
15+B:30-47 top500_models/*.pdb
```

```
rna_calc_inf.py -t 6n2v_with_FRETsequence_renum.pdb --target-
selection A:2-24+A:28-57+B:1-15+B:30-47 --model-selection A:2-
24+A:28-57+B:1-15+B:30-47 top500_models/*.pdb
```

The deformation index was calculated as $DI = RMSD / INF_{all}$.

FRET was predicted using donor and acceptor ACVs computed with *FRETraj* (39) as described in the main text.

### Supplementary Tables

**Supplementary Table 1.** Labeling chemistries for nucleic acids and fluorescent dyes currently implemented by *FRETlabel* (11, 40–43).

<table>
<tr><th>Position</th><th>Chemistry</th><th>Base</th><th>Dye</th></tr>
<tr><td rowspan="2">Internal</td><td>U/dT-C5</td><td>U<br>dT</td><td rowspan="5">sCy3<br>sCy5<br>Cy5.5<br>Cy7<br>Cy7.5<br>Alexa350<br>Alexa488<br>Alexa532<br>Alexa568<br>Alexa594<br>Alexa647<br>Atto390<br>Atto425<br>Atto465<br>Atto488<br>Atto495<br>Atto514<br>Atto520<br>Atto610</td></tr>
<tr><td>Etheno-adduct</td><td>A<br>C<br>dA<br>dC</td></tr>
<tr><td>5'-end</td><td>Phosphate</td><td rowspan="2">A<br>U<br>G<br>C<br>dA<br>dT<br>dG<br>dC</td></tr>
<tr><td rowspan="2">3'-end</td><td>Phosphate</td></tr>
<tr><td>Hydrazide</td><td>A<br>U<br>G<br>C</td></tr>
</table>

**Supplementary Table 2.** Dye and linker parameters used for ACV simulations.

| Parameter | symbol | value |
|---|---|---|
| Dye radii | $R_{1,\mathrm{Cy3}}$ (nm)<br>$R_{2,\mathrm{Cy3}}$ (nm)<br>$R_{3,\mathrm{Cy3}}$ (nm)<br>$R_{1,\mathrm{Cy5}}$ (nm)<br>$R_{2,\mathrm{Cy5}}$ (nm)<br>$R_{3,\mathrm{Cy5}}$ (nm) | 8<br>3<br>1.5<br>9.5<br>3<br>1.5 |
| Linker length<br>Linker width | $L_{\mathrm{linker}}$ (nm)<br>$W_{\mathrm{linker}}$ (nm) | 20.5<br>5 |
| Diameter of contact volume | $d_{\mathrm{CV}}$ | 3 |

**Supplementary Table 3.** *FRETraj* parameters used for calculating shot-noise broadened FRET efficiency histograms from MD simulations with explicit all-atom dyes or multi-ACVs.

| Category | Parameter | Symbol | Value |
|---|---|---|---|
| Dyes | Fluorescence lifetimes | $\tau_D$ (ns) | 1 |
| | | $\tau_A$ (ns) | 1.5 |
| | Quantum yields | $QY_D$ | 0.2 |
| | | $QY_A$ | 0.3 |
| | Angle between absorption and emission dipole | $\beta$ (°) | 10.5 |
| Bursts | Number of bursts | $n_{bursts}$ | 10000 |
| | Minimum photon threshold for analytical burst sizes | $N_{total,min}$ | 20 |
| | Maximum photon threshold for analytical burst sizes | $N_{total,max}$ | 150 |
| | Exponent of analytical burst size distribution | $\lambda$ | -2.3 |
| | Averaging regime | - | Ensemble |
| | QY correction | - | False |
| FRET | Förster radius | $R_0$ (nm) | 5.4 |
| | Dye quenching distance | $R_{quench}$ (nm) | 1.0 |
| | Gamma (compare simulation against gamma-corrected FRET experiment) | - | True |

### Supplementary Figures

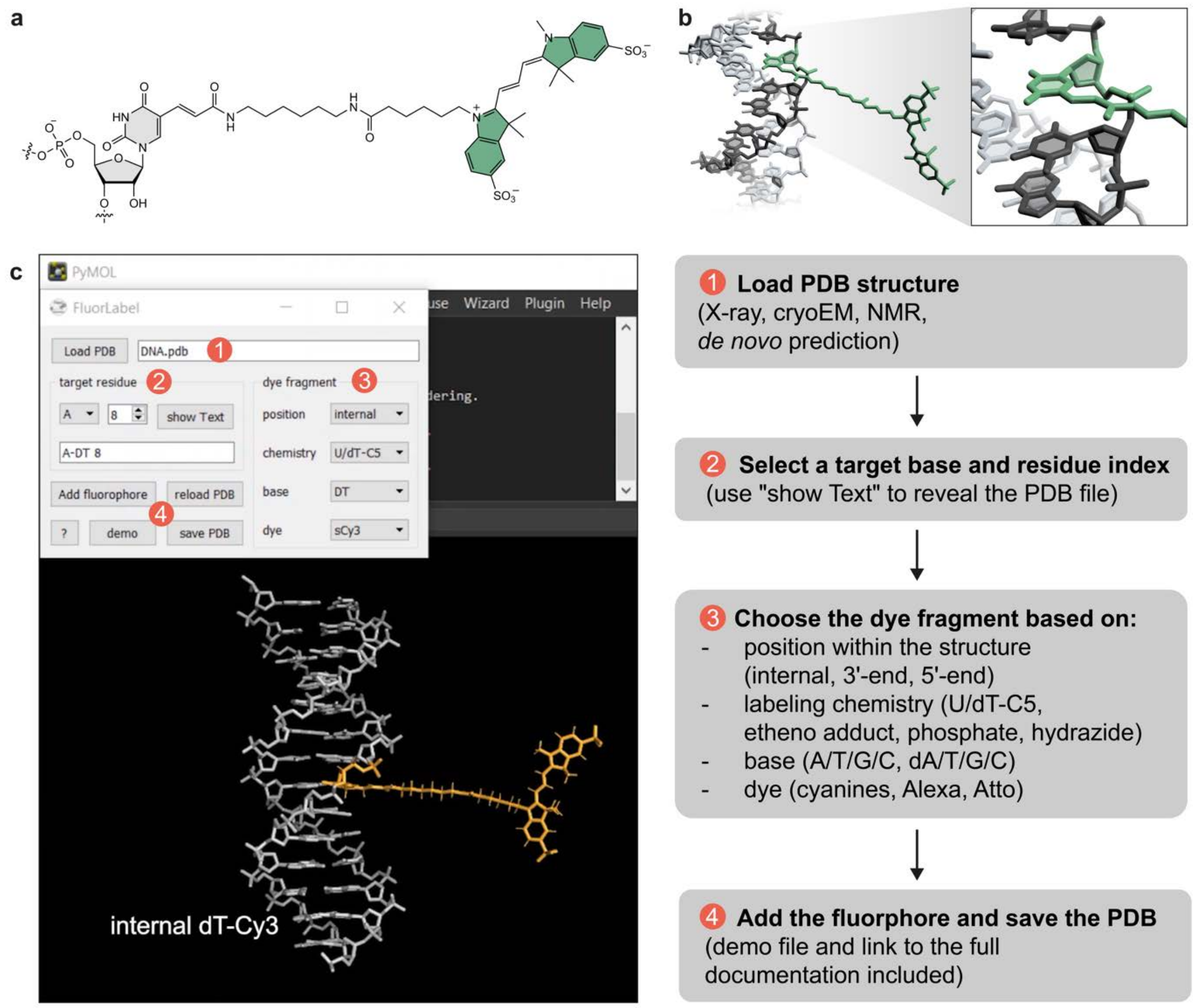


**Supplementary Fig. 1.** *In silico* fluorescence labeling of nucleic acids. **a** Chemical structure of a Cy3 fluorophore which is covalently attached to a C5-modified 2'-deoxyuridine. **b** Licorice representation of the Cy3 dye coupled to a DNA double helix. **c** Implementation of the automated *in silico* labeling as a PyMOL plugin. The user can select the target residue, the position (terminal, internal), the type of chemistry (e.g. dU-C5, etheno-adduct) and the dye (various cyanines, Alexa and Atto fluorophores).

**a**

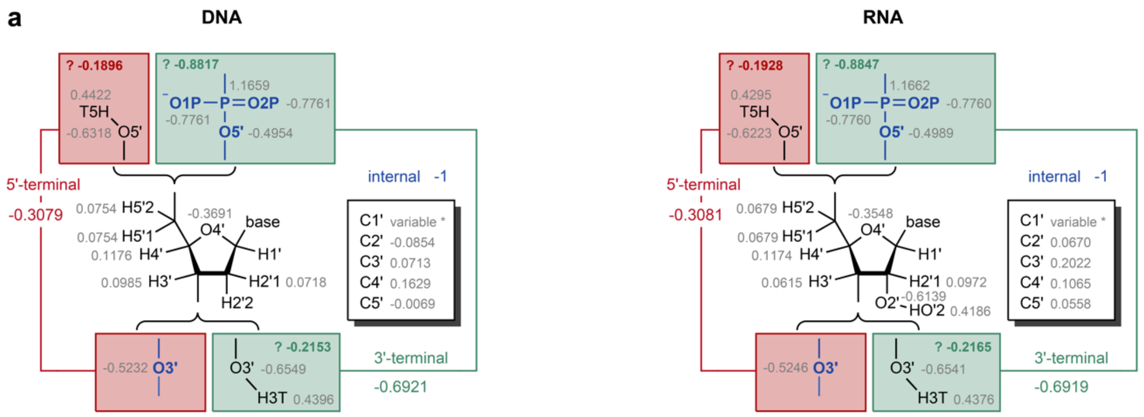


5'-terminal = -1 - (-0.8817) + (-0.1896) = -0.3079

3'-terminal = -1 - (-0.5232) + (-0.2153) = -0.6921

base + C1' + H1' = 0.1216

| | adenine | guanine | cytosine | thymine |
|---|---|---|---|---|
| C1' | 0.0431 | 0.0358 | -0.0116 | 0.0680 |
| H1' | 0.1838 | 0.1746 | 0.1963 | 0.1804 |

5'-terminal = -1 - (-0.8847) + (-0.1928) = -0.3081

3'-terminal = -1 - (-0.5246) + (-0.2165) = -0.6919

base + C1' + H1' = 0.1160

| | adenine | guanine | cytosine | uracil |
|---|---|---|---|---|
| C1' | 0.0394 | 0.0191 | 0.0066 | 0.0674 |
| H1' | 0.2007 | 0.2006 | 0.2029 | 0.1824 |

**b**

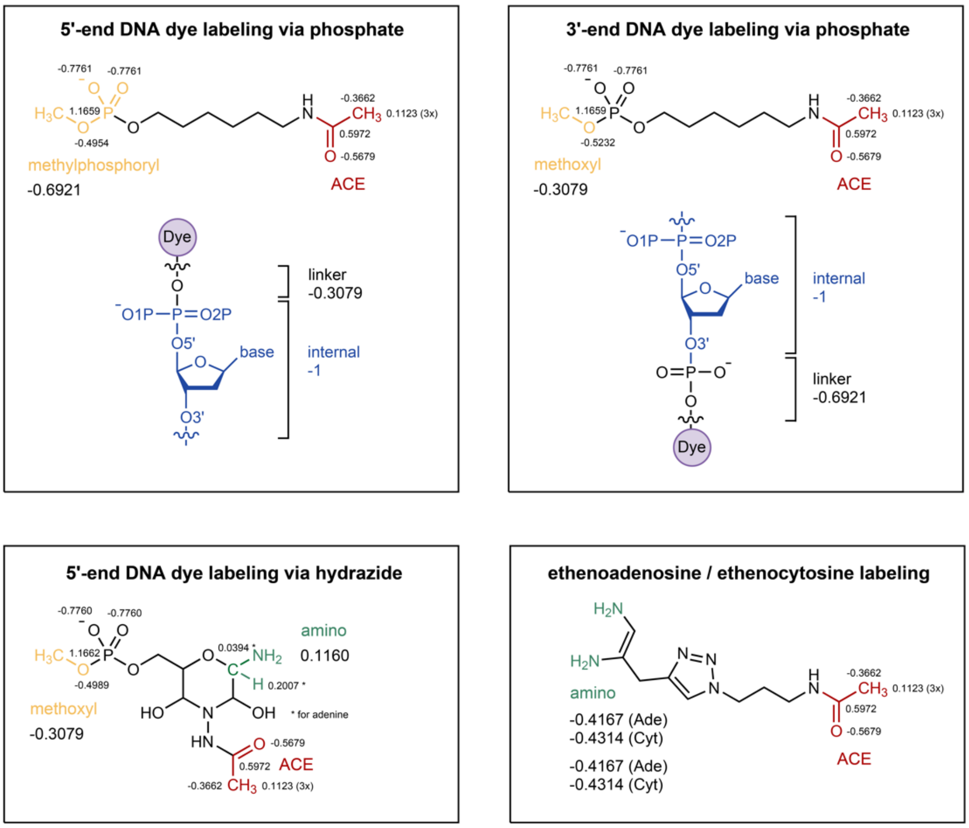


**Supplementary Fig. 2.** Force-field parameterization of linker atoms. **a** Partial charges of RNA and DNA residues in the AMBER force field derived by Cornell *et al*. (19) **b** Partial charges for the capping groups of the dye linker constrained during RESP fitting (16).

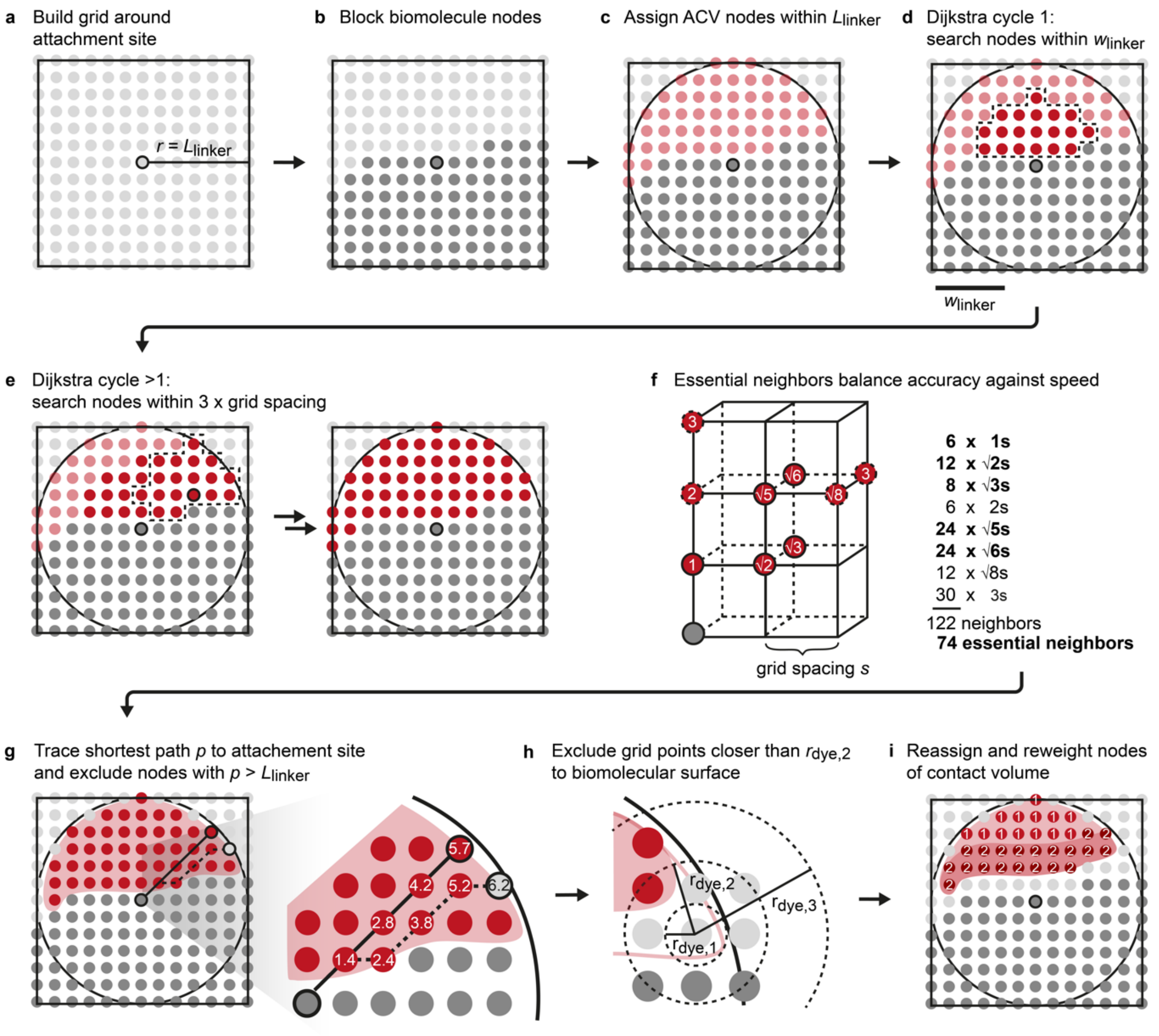


**Supplementary Fig. 3.** Simplified two-dimensional scheme of grid node assignment in the ACV algorithm. **a** A grid with radius $r = L_{linker}$ and grid spacing $s$ is constructed around the attachment site (black circle). **b** Grid nodes within the VdW surface of the biomolecule plus a clash radius $r_{clash}=w_{linker}/2$ are assigned to the biomolecule (dark gray spheres). **c** Grid nodes within $L_{linker}$ are tentatively allocated to the accessible volume (AV, light red spheres). **d** Dijkstra's algorithm is used to validate the ACV grid nodes, allowing only nodes with a shortest path $p > L_{linker}$ to the attachment site (see below). To avoid blocking the algorithm near the attachment site, all nodes within half of the linker width $w_{linker}$ are searched in the first round (nodes in dotted envelope). **e** In subsequent iterations, the grid is searched for nodes within a radius of 3s. **f** Hence, nearest neighbors can be located along a rank or on a diagonal but must always be closer than 3s from the current node. This results in a total of 122 neighbors. In practice, only 74 essential neighbors with distances 1s, √2s, √3s, √5s, √6s from the current node are added to the adjacency list to balance accuracy of the final volume against speed. **g** Each node is retraced to the attachment site and only nodes with a shortest path (i.e. cumulative sum) smaller than $L_{linker}$ are accepted. **h** Grid points located closer to the biomolecular surface than one of the dye radii (here middle radius $r_{dye,2}$) are excluded. **i** Nodes within $d_{CV}$ from the biomolecule's VdW surface are allocated to the contact volume and are assigned a weight according to Equation 3 (see main text).

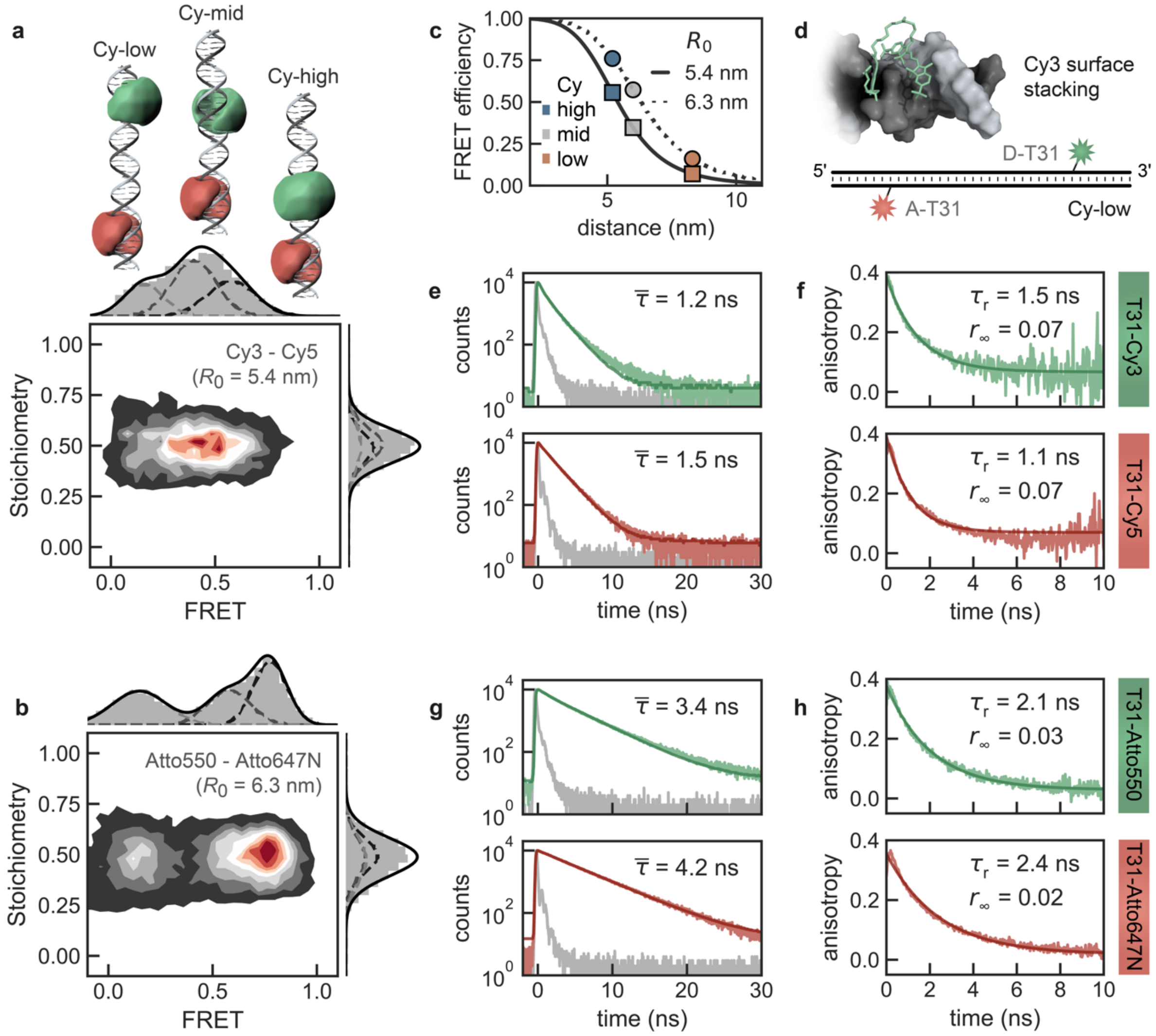


**Supplementary Fig. 4.** Comparison of different fluorophore pairs for FRET, fluorescence lifetime and anisotropy measurements on double stranded DNA. **a** 2D FRET-stoichiometry histogram of three cyanine labeled DNA double helices mixed in near equimolar amounts and recorded at 20 mM $Mg^{2+}$ and 5 mM NaCl at pH 7.5. Accessible volume pictograms indicate the different dye positions. **b** Control FRET measurements with Atto-dyes as used in the original benchmark study (10). **c** Expected transfer efficiencies for different Förster radii $R_0$ (Cy3-Cy5: 5.4 nm; Atto550-Atto647N: 6.3 nm) as a function of the inter-dye distance. **d** Surface interaction of Cy3 in the Cy-low sample used for lifetime and anisotropy experiments. In this duplex construct the donor fluorescence lifetime is least affected by FRET. **e/f** Lifetime and anisotropy decays of the cyanine dyes. **g/h** Same measurements as in e/f but for the Atto samples. The anisotropy decays are very similar to those of the cyanines although with an overall better signal-to-noise ratio due to the longer lifetimes of the Atto labels.

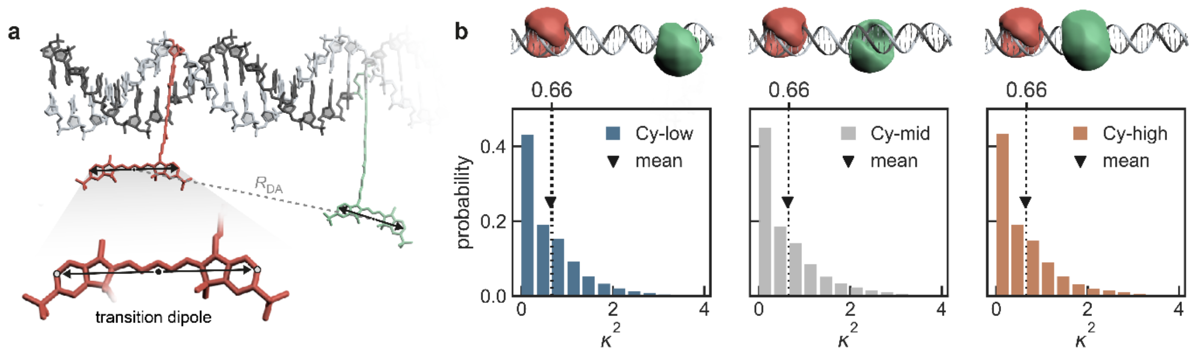


**Supplementary Fig. 5.** Fluorophores rotate isotropically in all-atom MD simulations. **a** Definition of the transition dipole by two peripheral carbon atoms in cyanine dyes. The donor acceptor distance $R_{DA}$ is given by the distance between the centers of mass of the two atoms on each dye. **b** $\kappa^2$-distributions of the three Cy-samples from all-atom MD simulations. The mean of the distributions coincides with the isotropic average ($\kappa^2$ = 2/3, dashed line) confirming that the ACV model, which presumes isotropic rotation of the fluorophores, is indeed applicable.

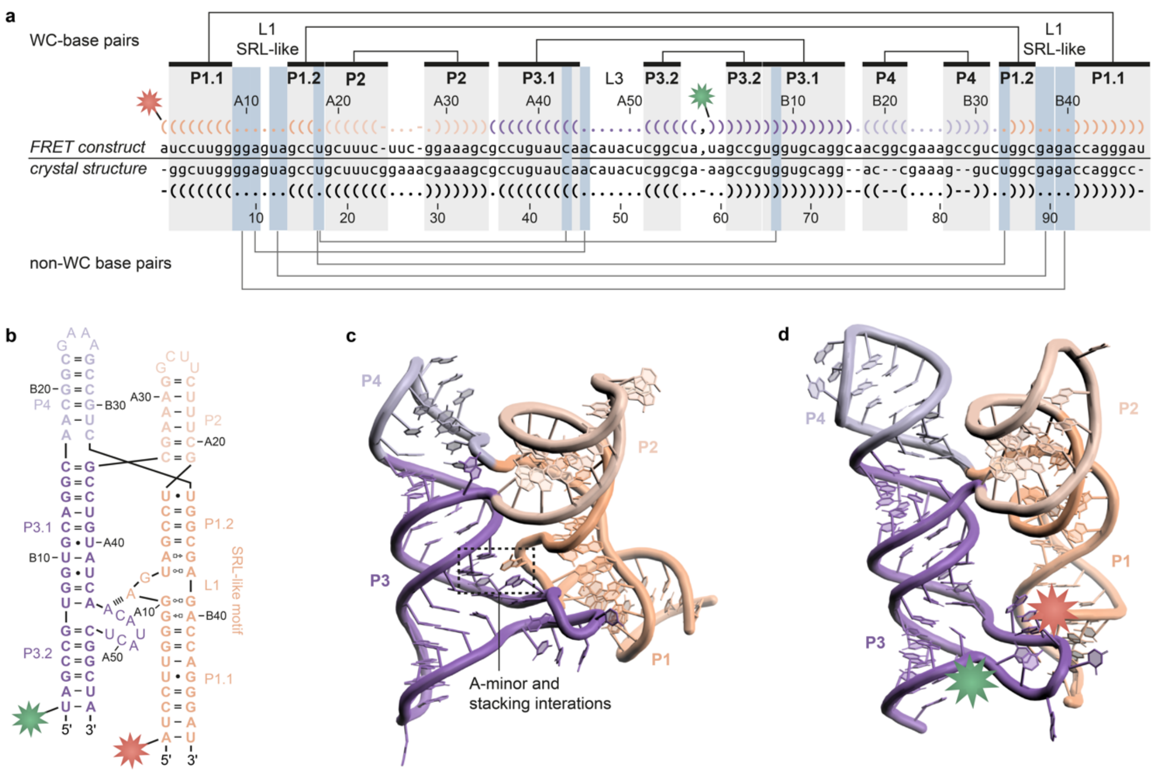


**Supplementary Fig. 6.** *De novo* structure prediction of the $Mn^{2+}$ riboswitch aptamer. **a** Sequence alignment of the *yybP-ykoY* RNA crystal structure (PDB: 6n2v, conformer 1) and the end-labeled FRET construct from Suddala et al. *Nat. Commun.* (2019) (34). Secondary structure and non-Watson-Crick (WC) tertiary contacts are annotated. **b** Secondary structure diagram of the FRET construct labeled at A1 (5'-end of chain A, Cy5) and U60 (5'-end of chain B, Cy3). L1 adopts a sarcin-ricin-like (SRL) motif harboring an extruding adenine which contacts L3 via an A-minor interaction. **c** Crystal structure of the riboswitch with the core forming base interactions indicated. **d** Best *de novo* structure from FRET-informed FARFAR2. While the core is incompletely packed, the global fold of the four-way junction is still recapitulated with an RMSD of 8.6 Å.
